# Computational Speed-Up of Large-Scale, Single-Cell Model Simulations Via a Fully-Integrated SBML-Based Format

**DOI:** 10.1101/2022.10.13.511603

**Authors:** Arnab Mutsuddy, Cemal Erdem, Jonah R. Huggins, Michael Salim, Daniel Cook, Nicole Hobbs, F. Alex Feltus, Marc R. Birtwistle

## Abstract

**Summary:** Large-scale and whole-cell modeling has multiple challenges, including scalable model building and module communication bottlenecks (e.g. between metabolism, gene expression, signaling, etc). We previously developed an open-source, scalable format for a large-scale mechanistic model of proliferation and death signaling dynamics, but communication bottlenecks between gene expression and protein biochemistry modules remained. Here, we developed two solutions to communication bottlenecks that speed up simulation by ~4-fold for hybrid stochastic-deterministic simulations and by over 100-fold for fully deterministic simulations.

**Availability and Implementation:** Source code is freely available at https://github.com/birtwistlelab/SPARCED/releases/tag/v1.1.0 implemented in python, and supported on Linux, Windows, and MacOS (via Docker).

**Contact:** Marc Birtwistle

**Supplementary information:** N/A

## Text

Recapitulating the behavior of single cells *in silico* is a grand challenge not only for systems biology, but also for biology in general. Such an accomplishment would imply that we have a thorough understanding of all the cellular and sub-cellular processes that give rise to relevant phenotypes. Such models could enable rational engineering for biotechnology applications, or forward predictions in precision medicine (Ahn-Horst *et al*., 2022; Mardinoglu *et al*., 2017; Spann *et al*., 2018; Uhlen *et al*., 2017). Large-scale and whole-cell modeling is a suitable foundation for meeting such challenges (Purcell *et al*., 2013; Macklin *et al*., 2014; Goldberg *et al*., 2018). The first such efforts focused on genome-scale metabolic modeling in multiple organisms (Patil and Nielsen, 2005; Thiele and Palsson, 2010). Subsequent efforts focused on integrating multiple “modules” in addition to metabolism (e.g. gene expression, signaling, etc.) in single-celled organisms such as *M. genitalium, E. coli, S. cerevisiae*, (Covert *et al*., 2008; Karr *et al*., 2012; Ye *et al*., 2020; Münzner *et al*., 2019) and a minimal lab-generated cell (Thornburg *et al*., 2022), but the lack of dedicated tools specifically for large-scale / whole-cell models presented roadblocks for reuse. Algorithmic developments included rule-based modeling to specify reactions more compactly (Faeder *et al*., 2009), and model composition tools, (Lopez *et al*., 2013; Gyori *et al*.,2017; Hoops *et al*., 2006; Somogyi *et al*., 2015) but large-scale models often still presented challenges. More recent work has provided such tools like AMICI that enables SBML-specified models to be simulated quickly, PEtab (Stapor *et al*., 2018; Schmiester *et al*., 2021) and Datanator (Roth *et al*., 2021) that specifies data formats for parameter estimation, formalisms that can help with unambiguous species naming, (Lang *et al*., 2020) and composition approaches such as ours that simplify model aggregation and expansion in ways that are compatible with efficient large-scale simulation algorithms and easy to reuse (Erdem *et al*., 2022). Not unexpectedly, however, there remains much work to be done to even technically enable large-scale and whole-cell modeling.

Here, we focused on improving communication between different modules as a major impediment for computation speed in large-scale modeling (**Fig. 1**). We used our recently published SPARCED model as a test case, a large-scale mechanistic model of proliferation and death signaling in single mammalian cells. This model consists of 141 genes, and 1196 unique biochemical species. It is built by translating a simple set of structured input text files into an SBML-compliant module that captures “protein biochemistry” (signaling) and is simulated using AMICI, and a module that captures “gene expression” using python. It can be simulated in a stochastic/deterministic mode, where gene expression dynamics follow Poisson-like processes, or a fully deterministic mode.

**Figure 1.**
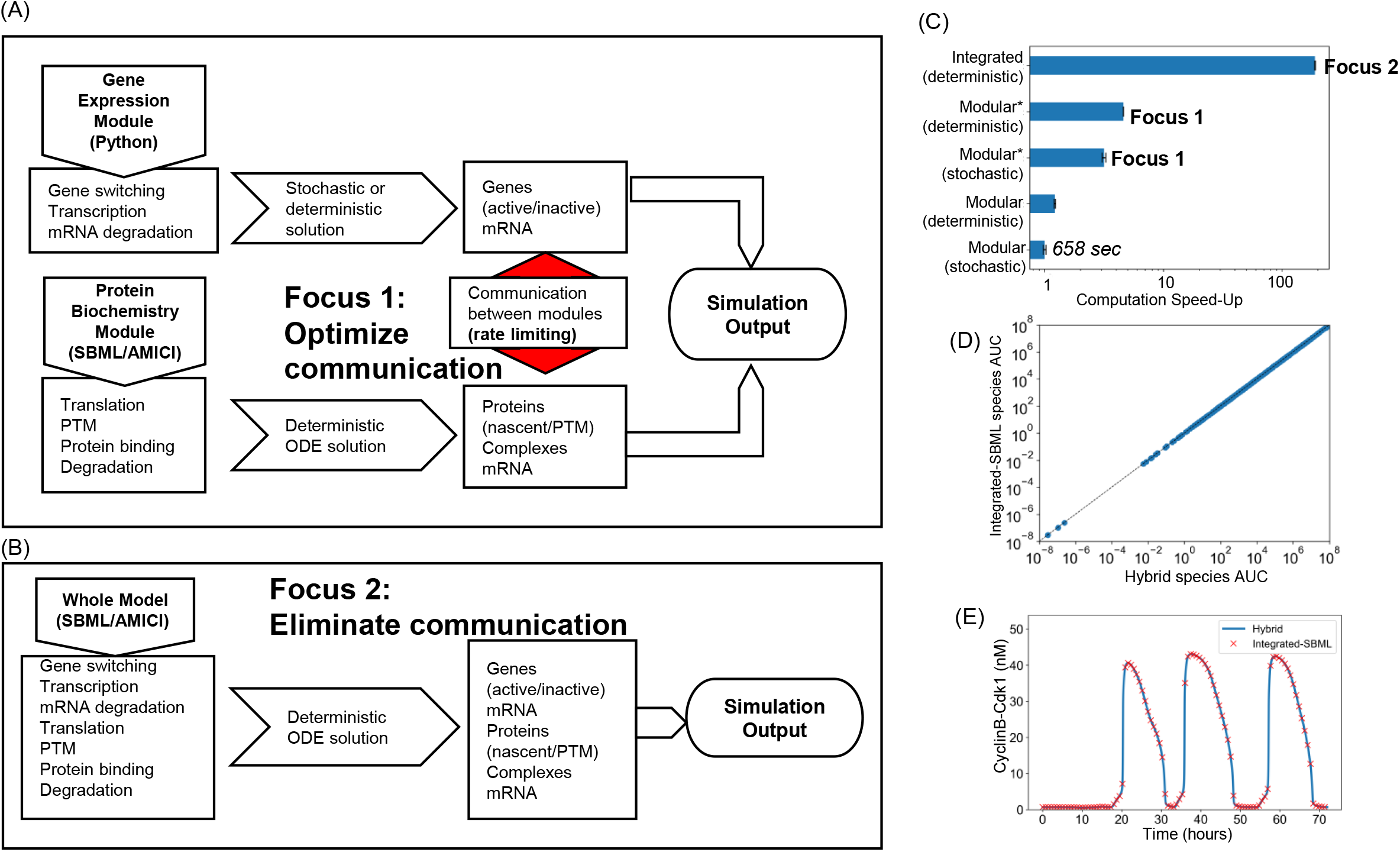
Computational speedup of the SPARCED model. **A.** Simulation workflow of the original SPARCED model highlighting the bottleneck of communication between the gene expression module and the protein biochemistry module. One speedup reported here targets that bottleneck for faster stochastic simulations. **B.** A new simulation workflow reported here that integrates the gene expression module with the protein biochemistry module using SBML, enabling large computational speed up for deterministic simulations. No solvers yet exist for stochastic simulations at this scale. **C.** Computation speed-up enabled by improving communication between modules (denoted by *; Focus 1) and by integrated SBML of the gene expression module with the protein biochemistry module (Integrated; Focus 2). Improving communication (Focus 1) yields 3-4 fold speed-up, and eliminating communication (Focus 2) yields ~100-fold speed-up. A relative speed of 1 corresponds to 658 seconds. Error bars are from 10 replicate simulations. Simulations were performed on Palmetto (Clemson’s HPC resource—Intel Xeon CPU 2.5 GHz). **D.** Area-under-curve for the dynamics of all model species in the original modular deterministic formulation and the integrated formulation. Simulated serum-starved MCF10A cells were treated with 1 nM EGF and 0.005 nM HGF and observed for 72 hours. **E.** An example trajectory for a biochemical correlate of cell division events from the simulations in D for both models, showing good agreement.

As is typical for large-scale models, communication between modules was done at specified simulation time steps, in our case every 30 simulated seconds. Using Python code profilers [cProfile, line_profiler], we identified communication between AMICI and python as rate limiting for simulation speed **(Fig. 1A—Focus 1).**Specifically, during each 30 second time step, results from the “protein biochemistry” module are saved in a “results” object defined within the AMICI library. However, accessing the state matrix via the Python object interface incurred expensive reconstructions of the full NumPy array from AMICI-managed memory. These overheads could be largely avoided, since only the last column of the state matrix (corresponding to the most recent timestep) was needed at each iteration. By using direct access to the SWIG pointer referencing these state variables, we were able to avoid re-reading state data, yielding a 34-fold simulation speedup **(Fig. 1C—Focus 1).**

However, we reasoned that a potentially better solution to improving module communication was to eliminate it altogether. This required a reformulation of the SPARCED model building scripts that translate the text input files into formats required for simulation, such that now a single SBML file was generated that can be simulated completely using AMICI (**Fig. 1B—Focus 2**). The drawback to this is that no efficient numerical solvers yet exist to perform stochastic simulations on such large models. Nevertheless, fully deterministic simulations are still of use in certain situations (e.g. during model initialization by which we convert the cellular context of the model using multi-omics data) (Bouhaddou *et al*., 2018; Barrette *et al*., 2018). After implementing this change, an over 100-fold computational speed-up was observed **(Fig. 1C— Focus 2).** We verified that simulation results obtained with this “integrated” model were identical to the original model to ensure that the reformulation of the model and its build process had not introduced any errors **(Fig. 1D-E).**

In conclusion, here we provide code that speeds up simulation of a large-scale model of cell behavior by ~4-fold for stochastic simulations and ~100-fold for deterministic simulations, by focusing on improving or eliminating communication between modules. We expect this to be impactful as a general strategy to further enable large-scale and whole-cell modeling, and also spur the development of simulation algorithms such as those that can perform stochastic simulations using an integrated formulation.

## Acknowledgements

This work was supported by the National Institutes for Health [R35GM141891 to M.R.B.].

## References

Ahn-Horst, T.A. et al. (2022) An expanded whole-cell model of E. coli links cellular physiology with mechanisms of growth rate control. npj Syst Biol Appl, 8, 1–21.

Barrette, A.M. et al. (2018) Integrating Transcriptomic Data with Mechanistic Systems Pharmacology Models for Virtual Drug Combination Trials. ACS Chem Neurosci, 9, 118–129.

Bouhaddou, M. et al. (2018) A mechanistic pan-cancer pathway model informed by multi-omics data interprets stochastic cell fate responses to drugs and mitogens. PLOS Computational Biology, 14, e1005985.

Covert, M.W. et al. (2008) Integrating metabolic, transcriptional regulatory and signal transduction models in Escherichia coli. Bioinformatics, 24, 2044–2050.

Erdem, C. et al. (2022) A scalable, open-source implementation of a large-scale mechanistic model for single cell proliferation and death signaling. Nat Commun, 13, 3555.

Faeder, J.R. et al. (2009) Rule-based modeling of biochemical systems with BioNetGen. In, Systems biology. Springer, pp. 113–167.

Goldberg, A.P. et al. (2018) Emerging whole-cell modeling principles and methods. Curr Opin Biotechnol, 51, 97–102.

Gyori, B.M. et al. (2017) From word models to executable models of signaling networks using automated assembly. Mol Syst Biol, 13, 954.

Hoops, S. et al. (2006) COPASI—a COmplex PAthway SImulator. Bioinformatics, 22, 3067–3074.

Karr, J.R. et al. (2012) A Whole-Cell Computational Model Predicts Phenotype from Genotype. Cell, 150, 389–401.

Lang, P.F. et al. (2020) BpForms and BcForms: a toolkit for concretely describing non-canonical polymers and complexes to facilitate global biochemical networks. Genome Biology, 21, 117.

Lopez, C.F. et al. (2013) Programming biological models in Python using PySB. Mol Syst Biol, 9, 646.

Macklin, D.N. et al. (2014) The future of whole-cell modeling. Current Opinion in Biotechnology, 28, 111–115.

Mardinoglu, A. et al. (2017) Personal model-assisted identification of NAD+ and glutathione metabolism as intervention target in NAFLD. Mol Syst Biol, 13, 916.

Münzner, U. et al. (2019) A comprehensive, mechanistically detailed, and executable model of the cell division cycle in Saccharomyces cerevisiae. Nat Commun, 10, 1308.

Patil, K.R. and Nielsen, J. (2005) Uncovering transcriptional regulation of metabolism by using metabolic network topology. Proceedings of the National Academy of Sciences, 102, 2685–2689.

Purcell, O. et al. (2013) Towards a whole-cell modeling approach for synthetic biology. Chaos, 23, 025112.

Roth, Y.D. et al. (2021) Datanator: an integrated database of molecular data for quantitatively modeling cellular behavior. Nucleic Acids Research, 49, D516–D522.

Schmiester, L. et al. (2021) PEtab—Interoperable specification of parameter estimation problems in systems biology. PLOS Computational Biology, 17, e1008646.

Somogyi, E.T. et al. (2015) libRoadRunner: a high performance SBML simulation and analysis library. Bioinformatics, 31, 3315–3321.

Spann, R. et al. (2018) A probabilistic model-based soft sensor to monitor lactic acid bacteria fermentations. Biochemical Engineering Journal, 135, 49–60.

Stapor, P. et al. (2018) Optimization and profile calculation of ODE models using second order adjoint sensitivity analysis. Bioinformatics, 34, i151–i159.

Thiele, I. and Palsson, B.Ø. (2010) A protocol for generating a high-quality genome-scale metabolic reconstruction. Nat Protoc, 5, 93–121.

Thornburg, Z.R. et al. (2022) Fundamental behaviors emerge from simulations of a living minimal cell. Cell, 185, 345–360.e28.

Uhlen, M. et al. (2017) A pathology atlas of the human cancer transcriptome. Science, 357, eaan2507.

Ye, C. et al. (2020) Comprehensive understanding of Saccharomyces cerevisiae phenotypes with whole-cell model WM_S288C. Biotechnology and Bioengineering, 117, 1562–1574.

